# Human retrosplenial theta and alpha modulation in active spatial navigation

**DOI:** 10.1101/2020.12.01.406124

**Authors:** Tien-Thong Nguyen Do, Chin-Teng Lin, Klaus Gramann

## Abstract

Spatial navigation is a complex cognitive process based on multiple senses that are integrated and processed by a wide network of brain areas. Previous studies have revealed the retrosplenial complex (RSC) to be modulated in a task-related manner during navigation. However, these studies restricted participants’ movement to stationary setups, which might have impacted heading computations due to the absence of vestibular and proprioceptive inputs. Here, we investigated neural dynamics of RSC in an active spatial navigation task where participants actively ambulated from one location to several other points while the position of a landmark and the starting location were updated. The results revealed theta power in the RSC to be pronounced during heading changes but not during translational movements, indicating that physical rotations induce human RSC theta activity. This finding provides a potential evidence of head-direction computation in RSC in healthy humans during active spatial navigation.

## Introduction

Spatial navigation is an essential human skill that helps individuals track their changes in position and orientation by integrating self-motion cues from linear and angular motion ***(McNaughton et al., 2006)**.* Spatial navigation involves several brain regions for spatial information processing ***(Doeller et al., 2010;Ekstrom et al., 2003)***, including the retrosplenial complex (RSC) ***(Epstein, 2008)***,to translate spatial representations based on egocentric and allocentric reference frames ***(Vann et al., 2009)***. Head direction (HD) cells that compute HD and orientation ***(Clark et al., 2010)*** provide vital information for the translation of information based on distinctive spatial reference frames. HD cells have been found in several brain regions, including the parahippocampal ***(Bellmund et al., 2016;Vass and Epstein, 2017)*** and entorhinal regions ***(Chadwick et al., 2015)*** as well as the thalamus ***(Shine et al., 2016)*** and the RSC ***(Baumann and Mattingley, 2010;Marchette et al., 2014;Shine et al., 2016)***. Theta oscillations have been described as an essential frequency underlying the computation of HD and spatial coding in grid cell models ***(Brandon et al., 2011;Koenig et al., 2011;Maidenbaum et al., 2018;Winter et al., 2015)*** in actively orienting rodents. Due to its anatomical connections, the RSC is also a central hub in a human brain network underlying several cognitive functions, including spatial orientation. The RSC has a direct connection to V4 (occipital), the pari-etal cortex, and the hippocampus and indirect connections to the middle prefrontal cortex ***(Vann et al., 2009)***. HD cells in the RSC encode both local and global landmarks simultaneously ***(Vann et al., 2009)**,* supporting the central role of the RSC in encoding and translating different spatial representations.

Several brain imaging studies using electroencephalography (EEG) to investigate the fast-paced time course of the neural basis of spatial cognitive processes have shown that the RSC translates between egocentric and allocentric spatial information ***(Gramann et al., 2010;Lin et al., 2015)***. The RSC works with the occipital and parietal cortices to translate egocentric visual-spatial information embedded in an egocentric (retinotopic) reference frame into an allocentric reference frame (***Vann et al., 2009***;***Lin et al., 2017)***. Most previous studies, however, were conducted in a stationary setup, and they did not investigate the neural mechanisms contributing to navigation in real-world environments, including motor efference, visual, proprioception, vestibular, and kinesthetic system information input or subject-driven allocation of attention ***(Cullen and Taube, 2017;Gramann, 2013)***. During active navigation, proprioceptive and motor-related signals significantly contribute to the estimation of self-motion, leading to higher accuracy in estimating travel distance and self-velocity ***(Becker et al., 2002;Frissen et al., 2011**; **Jürgens and Becker, 2006**; **Telford et al.,1995)***. Importantly, heading changes in naturalistic navigation are associated with vestibular input, which, together with the visual system and proprioception, is the driving input for HD cells ***(Cullen and Taube, 2017;McNaughton et al., 2006;Stackman et al., 2003)***. Although a few studies examined active spatial navigation in humans, their experimental designs did not reflect the brain dynamics associated with unrestricted near-real-life navigation experiences ***(Ehinger et al., 2014;Lin et al., 2015)***, or they relied on specific patient populations ***(Bohbot et al., 2017)***. In summary, there is little knowledge about the brain dynamics underlying spatial navigation in actively navigating human participants and how these dynamics subserve the computation and translation of spatial information embedded in distinct frames of reference for orientation.

In the present study, we investigated the brain dynamics of healthy human participants during active navigation. In an effort to overcome the restrictions of established imaging modalities, we adapted the Mobile Brain/Body Imaging (MoBI) approach ***(Gramann et al., 2011**,**2014;Makeig et al., 2009)**,* allowing physical movement of the participants. Thus, we recorded high-density EEG synchronized to head-mounted virtual reality (VR) while participants physically performed a spatial navigation task. Participants tracked their location and orientation by using self-motion cues from the visual, vestibular, proprioception, and kinesthetic systems as well as motor efferences. At the end of the trial, after traversing paths that included several turns and straight segments, participants pointed to previously encoded landmark locations. Their brain dynamics were analysed using independent component analyses (ICA) on high-density EEG data and subsequent source reconstruction. This approach allowed us to assess the brain dynamics originating in or near the RSC during the active navigation, focusing on i) the effect of active locomotion on brain dynamics in participants preferentially using an egocentric reference frame or an allocentric reference frame for navigation ***(Gramann et al., 2005)*** and ii) how multisensory convergence during the active movement changes the use of reference frames underlying active navigation compared with established desktop setups. The results demonstrate that active movement through space significantly changes the preferred use of spatial reference frames. Furthermore, naturalistic navigation reveals strong theta synchronization in the RSC during navigation phases that require heading computation and a substantial covariation of alpha desynchronization with the accuracy in a landmark pointing task.

## Materials and Methods

### Participants

Eighteen healthy adults (age 27.8±4.2, 2 females) participated in this experiment. All participants reported normal or corrected-to-normal vision. Each received $60 for their participation. The pro-tocol was approved by the University of Technology Sydney (UTS) (grant number: UTS HREC REF NO. ETH17-2095).

### Experiment design and tasks

#### Reference Frame Proclivity Test (RFPT)

Prior to the main experiment, the participants completed an online RFPT ***(Goeke et al., 2015;Gramann et al., 2005)*** to classify them as allocentric, egocentric, or mixed-strategy navigators. In the test, participants had to navigate through a tunnel on the flat screen monitor that included direction changes of various angles in the horizontal plane. When they reached the end, they were asked to select which of two homing vectors pointed back to the start of the tunnel. This choice, made over 40 trials, determined their navigation style: egocentric or allocentric if they consistently used that reference frame in at least 70%of the trials, or mixed-strategy navigators if they switched between frames. Of the 18 participants, five were egocentric navigators, seven were allocentric, and six were mixed.

#### Main Experiment Design

The experiment comprised a series of straightforward physical navigation exercises interspersed with spatial encoding/retrieval tasks but complicated by letter encoding/retrieval tasks to impose an additional cognitive workload on the participants. Each participant first performed four learning trials to familiarize themselves with the tasks and instructions and subsequently completed 23 sessions, with each session consisting of four trials, over the course of the full experiment. Each trial proceeded as follows. First, participants were shown a global landmark and given 4 seconds to remember its position before it disappeared. Participants were then given the following instruction: “Point to the landmark location”. They responded by pointing a controller at their reckoned location and clicking the hair-trigger (R1, figure 1A). A beep sounded to indicate their response had been registered and that they that should now start the first navigation task - straight walking with two to three direction changes over the walk while keeping track of their location in the space. More specifically, participants walked forward toward a floating red sphere at eye height, which disappeared once they reached it. A new red sphere then appeared, showing the next direction and distance and disappearing once they reached that, and so on. In figure 1A, this task is denoted as “walk 1x”, where 1 indicates the first walk and x indicates the number of red spheres. Once participants reached the last red sphere, they stopped walking and the text “Attention” appeared in front of them for 3 seconds, signaling the first spatial retrieval task was about to begin. First, participants were instructed to “Point to the landmark location” by pointing their controller to the landmark location as they remembered it and clicking the hair-trigger (R2a, figure 1A). Next, two homing arrows appeared in front of them - one pointing left, the other right - and they were asked: “Where is the starting location?”. Responses were given by pointing their controller at one of the arrows and clicking the trigger (R2b, figure 1A). The letter encoding task to impose additional cognitive burden followed. The participants were shown a series of 3, 5, or 7 letters of the English alphabet at one-second intervals between letters and asked to remember them. The number of letters chosen and the order in which the letters appeared were both random. Three levels of difficulty were included to avoid familiarity with the task, ensuring the cognitive load remained high. Then, three seconds after the last letter appeared, participants were shown a random letter and asked whether or not that letter belonged to their letter list. Clicking the trigger indicated yes; pressing the touchpad indicated no (R3, figure 1A). Participants then had 2 seconds of rest before starting the second walk. A beep sound signaled the beginning of the second navigation task (walk 2x, figure 1A), which followed the same straight walk to the red sphere format as the first. However, this time, the participants had to remember their letter list as well as keep track of their orientation toward the landmark location and starting position. When the second walk was finished, the participants were asked to do three things: first, to confirm whether or not a random letter belonged to their letter list (R4, figure 1A); second, to point to the landmark location; and, third, to point to the starting location (R5a and R5b, figure 1A). The next trial started when the participant indicated their readiness by clicking both bottom grips on the controller. The maths of four trials with 2 to 3 random turns in each of two walks meant that each session consisted of a total of 20 turns, as shown in figure 1B. After 20 turns, the participant reset his or her location to dot number 0 before starting the next session to ensure that the total navigation segments were within the experimental space. The RFPT in the active walk condition was assessed only at point 12 (figure 1B) based on the participant’s response to the left arrow (egocentric) or right arrow (allocentric).

**Figure 1.**
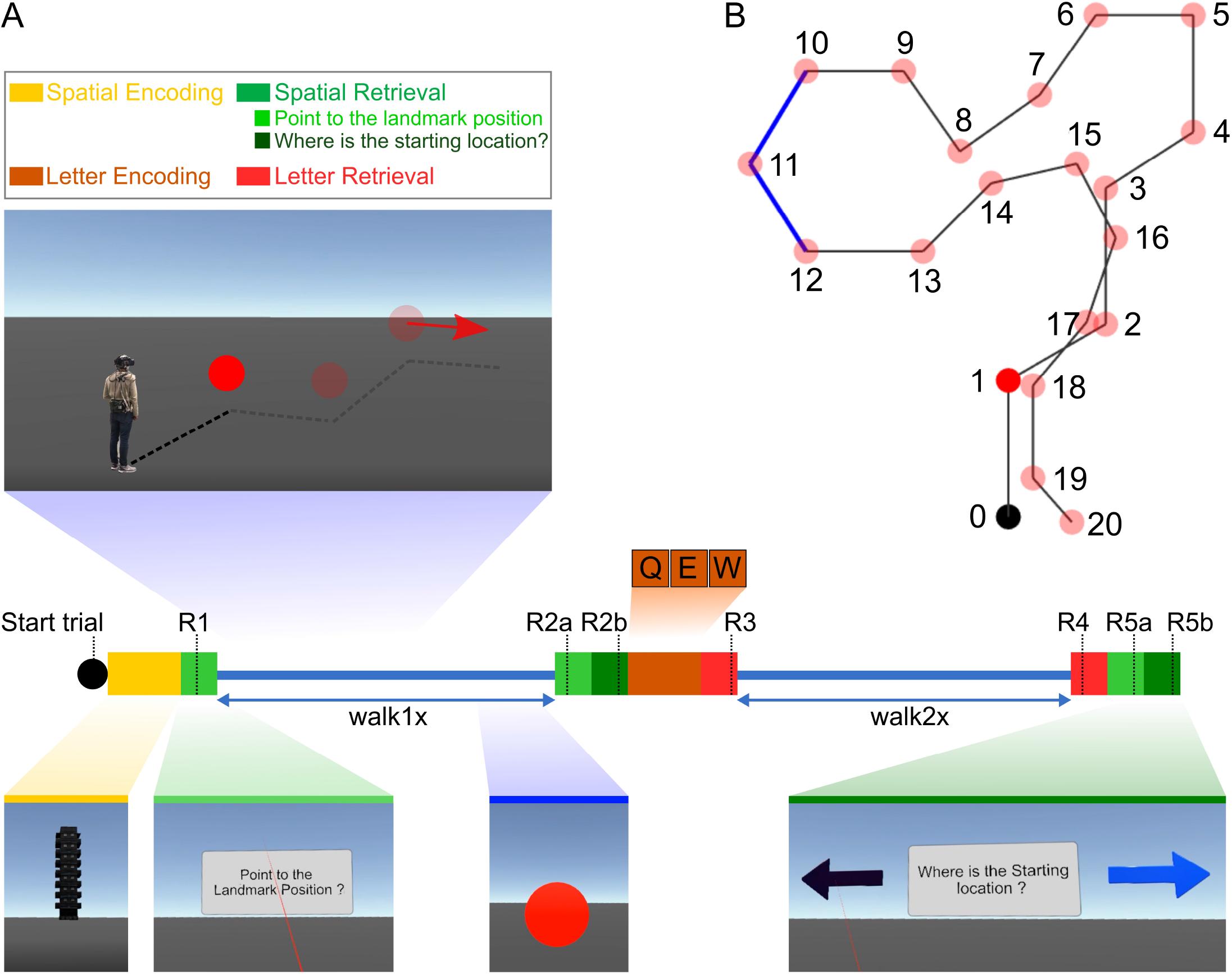
Experimental design. (**A**) At the beginning of the trial, the participants were given 4 seconds to remember a landmark position that appeared approximately 200 meters in front of them. They then performed two physical navigation tasks, walk1× and walk2×, each of which contained 2 to 3 random turns. After walk 1, they were also asked to remember a series of 3, 5, or 7 letters of the English alphabet (the number of letters chosen and the order in which the letters appeared were both random). Between the two walks and at the end of each trial, the participants were asked to point to the landmark location and their starting location and to confirm whether a randomly chosen letter appeared in the list of letters they had been asked to remember. Green squares indicate the landmark pointing task (R1, R2a, and R5a) and the starting point task (R2b and R5b). Red squares indicate the letter retrieval task. (**B**) Full participation in the experiment constituted 23 sessions, where each session consisted of four trials, each trial consisted of two walks, and each walk contained 2 to 3 turns in random directions. As such, each session involved a total of 20 turning points, shown as red dots. After 20 turns, the participant reset his or her location to dot number 0 before starting the next session to ensure that the total navigation segments stayed within the experimental space. The reference frame proclivity test (RFPT) was based on the participant’s response at dot number 12, where the answers for homing directions were clearly distinguishable between two the RFP strategies. Participants were considered egocentric or allocentric if their response was the left arrow or right arrow, respectively.

#### Data Recordings

The scenario was developed in Unity (version 2017.3) with the VRTK plug-in and performed in a VR environment using a head-mounted display (HTC Vive Pro; 2x 1440 x 1600 resolution, 90 Hz refresh rate, 110° field of view). All data streams from the EEG cap, and head-mounted display were synchronized by the Lab Streaming Layer ***(Kothe, 2014)***. The EEG data were recorded from 64 active electrodes placed equidistantly on an elastic cap (EASYCAP, Herrsching, Germany) with a sampling rate of 500 Hz (LiveAmps System, Brain Products, Gilching, Germany). The data were referenced to the electrode located closest to the standard position, FCz. The impedance of all sensors was kept below 5 kΩ.

### EEG analysis

#### Pre-processing

All pre-processing steps were performed using MABLAB version 2018a (Mathworks Inc., Natick, Massachusetts, USA) and custom scripts based on EEGLAB version 14.1.2 ***(Delorme and Makeig, 2004)*** (supplementary figure 3.1). The raw EEG data were downsampled to 250 Hz before applying a bandpass filter (1-100 Hz). Line noise (50 Hz) and associated harmonics were removed using the cleanline function in EEGLAB. Dead channels were then removed based on flatline periods (threshold=3 seconds), correlations with other channels (threshold=0.85), and abnormal data distributions (standard deviation=4). Missing channels were interpolated by spherical splines before re-referencing to the average of all channels. Noisy portions of continuous data were removed through automatic continuous data cleaning based on the spectrum value (threshold=10 dB) and power with a criteria of maximum bad channels (maximum fraction of bad channels=0.15) and relative to a robust estimate of the clean EEG power distribution in the channel ([minimum, maximum]=[-3.5 5]). Then, adaptive mixed independence component analysis (AMICA) ***(Delorme et al., 2012;Palmer et al., 2012)*** was used to decompose the data into a series of statistically maximally independent components (ICs) with the time source as the unit. The approximate source location of each IC was computed using the equivalent dipole models in EEGLAB’s DIFIT2 routines ***(Oostenveld and Oostendorp, 2002)***. Last, the spatial filter matrix and dipole models were copied back to the pre-processed but uncleaned EEG data (no cleaning in the time domain) for further analysis (supplementary figure 3.1).

#### Event-Related Spectral Perturbation (ERSP)

Epoched data sets for each walking condition (walk1× or walk2×)were extracted at the onset of navigation for a length of 14.5 seconds, including a baseline period of 2.5 seconds prior. Bad epochs were identified and removed in the sensor space (for strongly affected head and body motion artifacts) and subsequently in the component space (for artifact noise in the ICs). (i) In the sensor space, the bad epochs were labeled based on the epoch mean, standard deviation (std=5), absolute raw value (threshold value=1000 μV), and kurtosis activity (refer to the pop_autorej.m function in EEGLAB). From 2714 epochs, 12.5%were deemed bad and removed. (ii) In the component space, the time-warped ERSP patterns for each epoch were calculated by computing single spectrograms for each IC and single trial based on the newtimef() function using the standard parameters. The timewarp option was used to linearly warp each epoch to a standard length based on the median time point of sphere collision events. The time period before the onset of the active navigation at the beginning of the trial was used as the baseline to estimate the ERSP for each epoch with divisive baseline correction ***(Grandchamp and Delorme, 2011)*** to minimize single-trial noise. Epochs were then identified as bad if the z-score of the ERSP epoch was larger than 3 standard deviations from the median of all ERSP epochs (see the power panels in (figure 2, figure 5, and figure 4) Approximately 6%of the epochs were bad and removed based on their ERSP in the component space.

**Figure 2.**
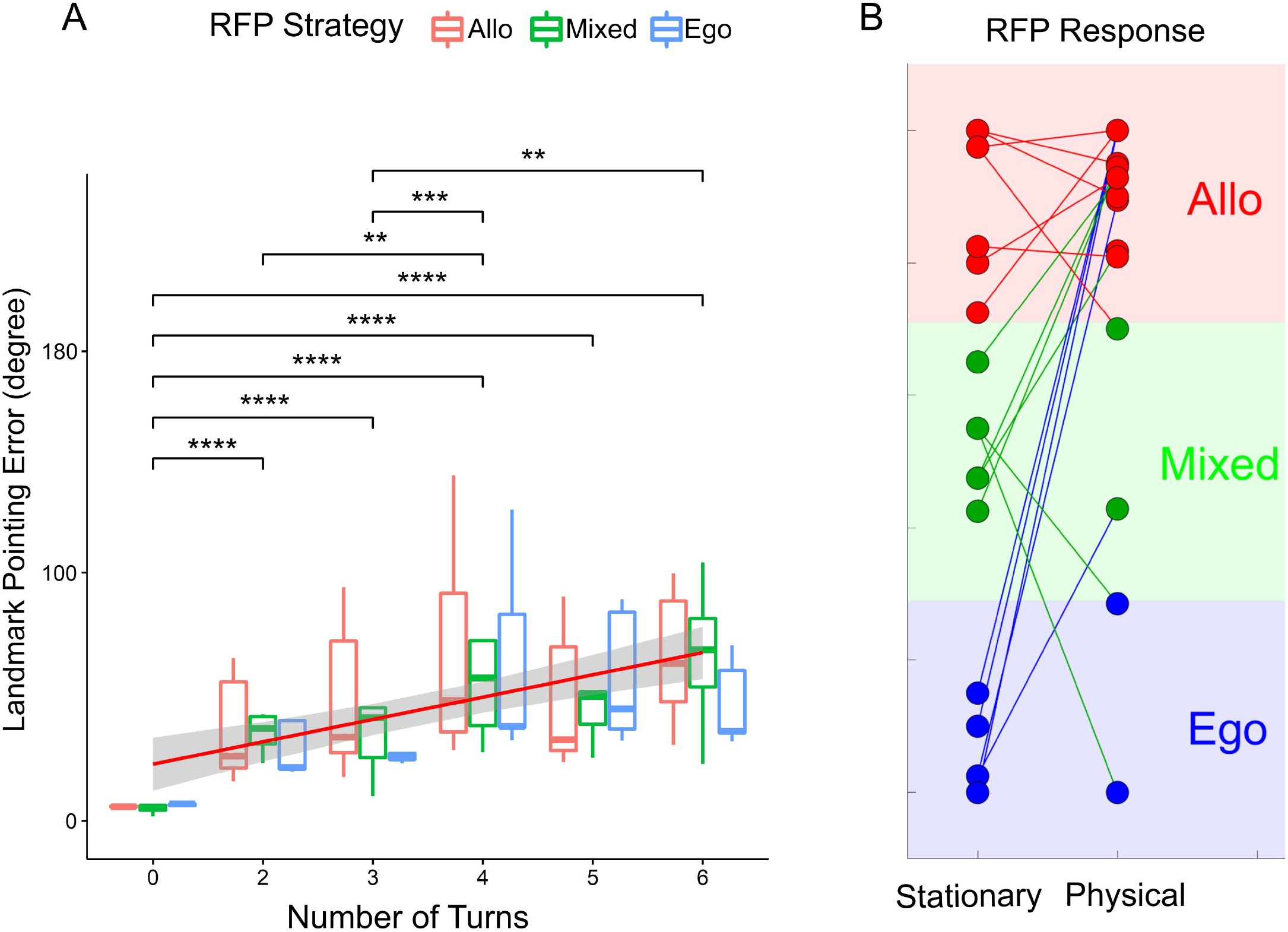
The results of participant behaviour. (**A**) Participant behavior in the landmark pointing task. The X axis indicates the number of turn points in the trial, the Y axis indicates the absolute error of the participants when they performed the landmark pointing task (the error was measured by the angular difference between the pointing vector and the participant to landmark vector). The regression was visualized by the red line (*, **, ***, **** indicated for p<.05, p<.01, p<.001, p<.0001 respectively, FDR-corrected). (**B**) The RFPT results were in both the passive condition (stationary test with the tunnel paradigm) and active condition (based on the participant’s response at position 12, path 3 in the Figure 1B). Three groups of strategies egocentric, mixed and allocentric were color coded in green, blue and red, respectively.

#### Clustering

We first clustered the ICs based on the conventional EEGLAB k-means method. Then, we repeated the clustering process 10000 times before performing an evaluation to identify the best cluster of interest (COI) based on the region of interest (ROI) cluster centroid ***(Gramann et al., 2018)***. All ICs with an RV of less than 30%for all participants were grouped based on their dipole locations (weight=6), ERSPs (weight=3), mean log spectra (weight=1), and scalp topography (weight=1). Then, the weighted IC measures were summed and compressed with principal component analysis (PCA), resulting in a 10-dimensional vector, followed by the k-mean method (with 25 clusters). The target cluster centroid in Talairach space (RSC, x=0, y=-45, z=10) was evaluated from 10000 clustering results based on the score of each cluster solution, including: (i) the number of participants (weight=2); (ii) ratio of the number of ICs per participant (weight=-3); (iii) cluster spreading (mean squared distance of each IC to the cluster centroid) (weight=-1); (iv) mean RV of the fitted dipoles (weight=-1); (v) distance of the cluster centroid to the ROI (weight=-3); and (vi) Mahalanobis distance to the median distribution of the 10000 solutions (weight=-1). The final COI score of-1.7 was derived from 15 participants, 29 ICs, a spread of 677, a mean residual variance (RV) of 11.76%, and a distance of 7.3 units in Talairach space. In the ERSP group-level analysis, the ERSP at COI was calculated first at the IC level, then at the participant level, and finally at the group level. The timefrequency data of all ICs of the same participant were averaged. Then, the ERSPs of all participants were averaged for the final ERSP at the group level. The statistical test for ERSPs was performed by a permutation test with 2000 permutations and a multiple comparison correction using the false discovery rate (FDR, *α*=0.05).

## Results

### Behavior

The navigation routes participants were asked to follow were pre-defined. First, they were shown a global landmark and asked to pointto it. Afterthe landmark disappeared, they followed a path with several turns and, after two or three changes in course, they were asked to point to the (now invisible) landmark. The navigation task resumed with two or three more turns, and the trial concluded by asking them to point to the invisible landmark a final time. The landmark pointing error (in degrees) was statistically significantly different at the different number of turns (NT) using the Friedman test, (*χ*^2^(5)=53.34, p<.0001). The pairwise Wilcoxon signed rank test (with false discovery rate-FDR corrected) between groups revealed statistically significant differences in landmark pointing error between NT0 and NT2 (p=.000046); NT0 and NT3 (p=.000046); NT0 and NT4(p=.000046); NT0 and NT5 (p=.000046); NT0 and NT6 (p=.000046); NT2 and NT4 (p=.007); NT3 and NT4 (p=.00011); NT3 and NT6 (p=.003). The median error was 5.79° (degrees) with 0 turns, 29.63° after 2 turns, 32.76° after 3 turns, 45.57° after 4 turns, 44.94° after 5 turns, and 60.99° after 6 turns.

### Event-related Spectral Perturbation (ERSP)

Repeated k-means clustering of independent components (ICs) resulted in 25 clusters with centroids located in several structures of the brain including the frontal cortex, the left and right motor cortices, the parietal cortex, the RSC, and the occipital cortex. Focusing on power spectrum changes in the RSC cluster, we computed event-related spectral perturbation (ERSP) in the frequency range of 3 to 50 Hz (figure 3). Broadband alpha and beta desynchronization were prominent during the straight segments of navigation, replicating previous results from both passive and active navigation studies ***(Gramann et al., 2010;Lin et al., 2015;Ehinger et al., 2014)***. In addition, a prominent theta burst became apparent directly after each turn, i.e., at the time when people were computing heading changes before proceeding along the next straight path. Moreover, the theta burst was present during all turns, while the alpha and low beta desynchronization became more desynchronized as the number of turns increased (supplementary figure 3.3).

**Figure 3.**
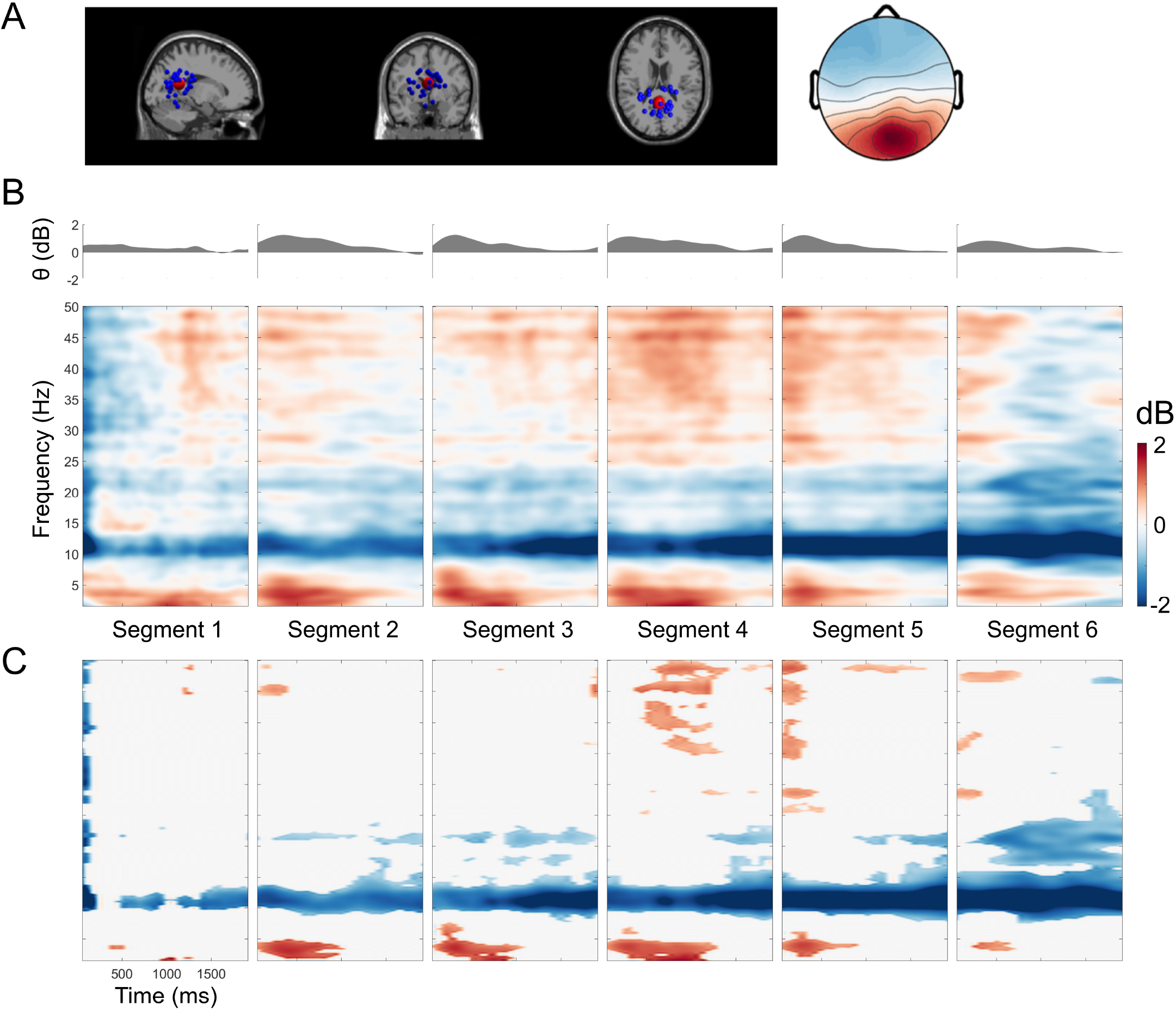
Retrosplenial complex (RSC) event-related spectral perturbation (ERSP). (**A**) Dipole locations of independent component (in or near the retrosplenial complex (RSC) cluster at the sagittal, coronal, and top view respectively and the corresponding mean of the scalp map. (**B**) The RSC ERSP with respect to the segment of turns from 1 to 6 turns. (**C**) The permutation test (n=2000, FDR-correlated) of the RSC ERSP in 6 segments.

**Figure 4.**
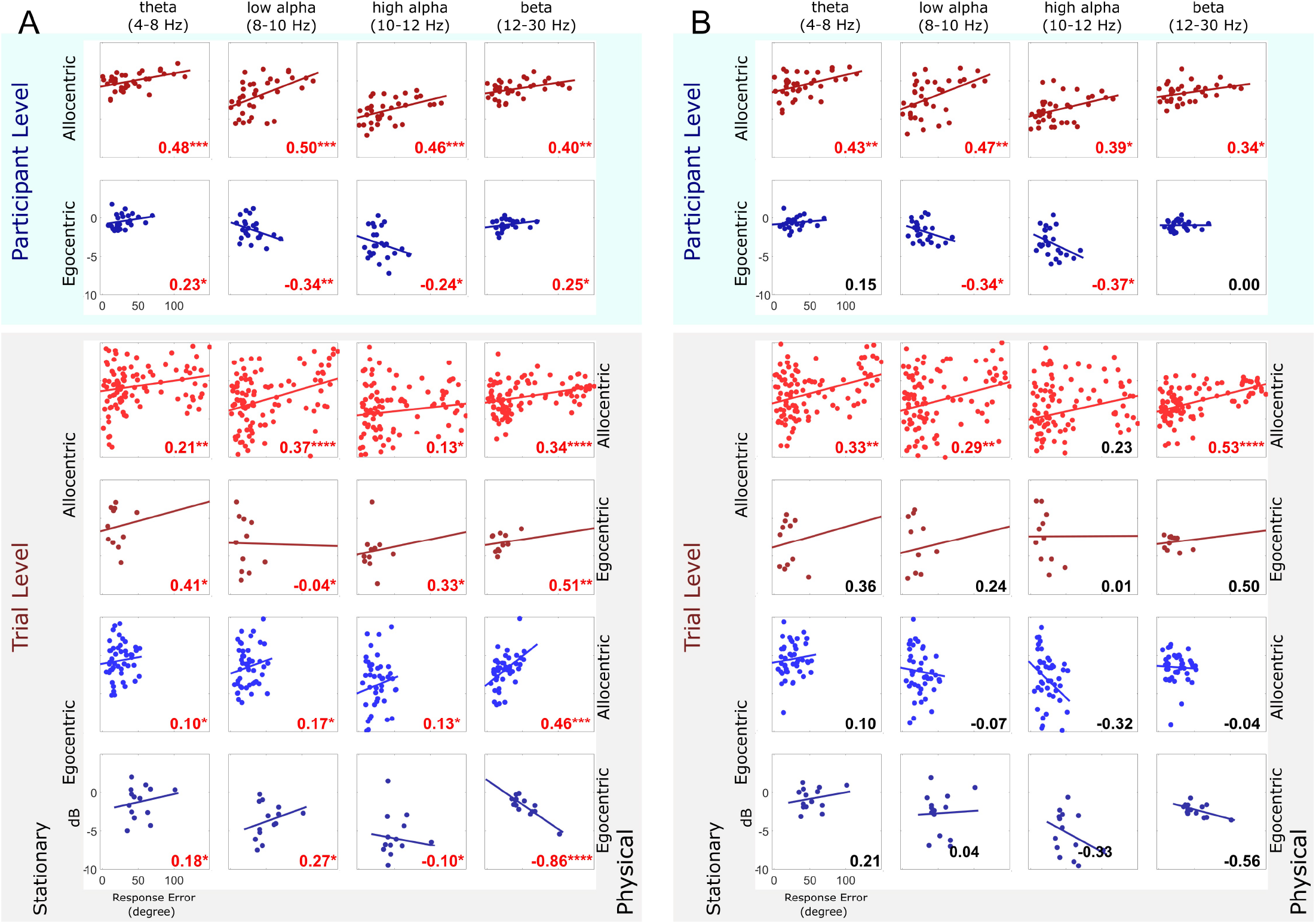
RSC ERSP correlated with participant performance in the first 10 percent (**A**) and mid 10 percent (**B**) of the segment length at both the participant and trial level respectively. The top figure shows the computed correlation coefficients (the number at the bottom right corner, red color indicated for statistical significance;*, **, ***, **** indicated for p<.05, p<.01, p<.001, p<.0001 respectively - FDR-corrected) between individual performance and RSC ERSP at participant level in different frequency band at theta (4-8 Hz), low Alpha (8-10 Hz), high Alpha (10-12 Hz), and beta (12-30 Hz). The row indicates the RFP strategy in the passive RFPT response: red indicates an allocentric, and blue indicates an egocentric strategy. The bottom figure shows the correlation coefficients computed between individual performance and RSC ERSP at the trial level in same the range of frequency as in the top figure. The color-coding indicates the RFP strategy;the lighter color in the same RFP strategy group indicates an active allocentric response, while the darker color indicates an active egocentric trial response.

### Neural Correlations with Spatial Updating

#### Heading computations

Next, we calculated the correlations between power modulations in the RSC and the landmark pointing errors. Fora comprehensive analysis, we divided the power modulations into different frequency bands and the pointing errors by allocentric or egocentric reference frames for the entire course of the navigation task. Further, to assess the impact of rotational compared with translational movement on RSC spectral modulations, we extracted the first 10%of each segment, which included the turn, and the middle 10%of each segment (50-60%), where participants were only moving in a straight line, and calculated the correlations between power in different bands and pointing errors with just these segments.

#### Participant-level Analysis

The allocentric group showed a significant positive correlation between errors in the landmark pointing tasks and power changes in the following broadband frequency ranges: theta (r(34)=0.48, p=.00048), low alpha (r(34)=0.5, p=.00051), high alpha (r(34)=0.46, p=.00047)and beta band (r(34)=0.4, p=.0012) (FDR-corrected) (see figure 4A). Egocentric participants demonstrated a negative correlation between the performance pointing task (error) and power changes in the following frequency ranges: low alpha (r(22)=−0.34, p=.0.006), and high alpha (r(22)=−0.24, p=.011)(FDR-corrected)(figure 4A). The inverse pattern of correlation coefficients of power and individual performance error for allocentric and egocentric participants was consistent across the entire navigation phase (10×10%time bins) (supplementary figure 4.1), revealing that RSC activity depends on their stationary reference frames used to encode and integrate the spatial information.

#### Trial-level Analysis

In contrast to previous stationary navigation studies ***(Lin et al., 2015;Ehinger et al., 2014)***, participants moved actively through the environment, receiving sensory feedback from not only the visual system but also converging sensory evidence about changes in position and orientation from their vestibular and proprioceptive systems. This phenomenon opens the possibility that the participants’ preferred use of spatial reference frames might change depending on the sensory input available to them ***(Gramann, 2013;Goeke et al., 2015)***. Therefore, we further investigated the relationship between individual performance error and power changes in RSC on a single-trial level. The pointing response at the end of the path required a binary decision regarding whether the homing location was located to the left or right with respect to the current position and orientation of the navigators (at point 12, figure 1B). The results of the single-trial reference frame classification demonstrated that a large portion of participants preferentially used an egocentric reference frame in stationary setups but switched to an allocentric reference frame with active navigation (figure 2B). In contrast, participants with a preference for using an allocentric reference frame in stationary setups kept the same allocentric reference frame in the active navigation scenario (figure 2B). Importantly, whenever participants used an allocentric reference frame in the pointing task, irrespective of their habitual proclivity toward an egocentric or an allocentric reference frame in stationary settings, there was a positive correlation between the pointing error and power changes in the lower alpha band (r(100)=0.37, p=.000037 for stationary allocentric; r(44)=0.17, p=.015 for stationary egocentric), higher alpha band (r(100)=0.13, p=.013 for stationary allocentric; r(44)=0.13, p=.016 for stationary egocentric), and beta band (r(100)=0.34, p=.00009 for stationary allocentric; r(44)=0.0012, p=.0.00015 for stationary egocentric). However, pointing based on an egocentric reference frame (in physical navigation) still revealed a negative covariation with power in the higher alpha band (r(12)=-0.10, p=.024) (dark-blue color, figure 4A) and the beta band (r(12)=-0.86, p=.000035 for stationary egocentric). Thus, the single-trial reference frame analyses clearly revealed a systematic and more pronounced desynchronization in the alpha band whenever an allocentric reference frame was used to respond to a homing challenge.

## Discussion

Spatial navigation is vital to purposeful movement as it requires a representation of our position and orientation in space as well as our homing trajectories. In this study, we explored these processes through a typical stationary navigation task but also a physical navigation task where the participants actually moved through a large virtual space while we recorded and analyzed their brain dynamics usinga MoBI approach ***(Makeig et al., 2009**; **Gramann et al., 2011**, **2014**; **Jungnickel et al., 2019)***. We found that participants with a proclivity for using an egocentric reference frame in stationary navigation tasks switched to an allocentric reference frame during physical navigation. In contrast, participants with an allocentric proclivity in stationary tasks still used the same reference frame during physical navigation. Importantly, using this modified MoBI approach provided this first-ever opportunity to describe theta synchronization in the RSC during heading computation in actively rotating navigators. From our analyses, we find that alpha desynchronization in the RSC occurs when retrieving spatial information from an allocentric reference frame and translating it into an egocentric location pointing response.

Remarkably, navigators switched from their preferred egocentric reference frame to an allocentric reference frame when they were allowed to actively move through space. Thus, the reference frame proclivity (RFP) observed in stationary navigation tasks is not consistent with that observed in active navigation tasks, including naturally occurring sensory feedback from the visual, vestibular, proprioception, and kinesthetic systems. More specifically, the majority of egocentric navigators switched to an allocentric reference frame during physical navigation, while the allocentric group consistently used their preferred allocentric strategy. To anchor a cognitive map with the environment, navigators can use local and global landmarks (e.g., a mailbox, a building) and/or self-motion cues. In our navigation scenario, we gave the participants a single prominent landmark only at the very beginning of a trial that was invisible for the rest of the navigation task. Participants then moved through space, walking straight toward a point and then locating and changing directions several times while moving away from their starting location (figure 1). Consequently, the participants tended to derive their orientations and positions in space by converging multiple sensory inputs to represent the original global landmark position. Most participants, including the egocentric strategy group, responded as allocentric navigators in the homing direction test (figure 2B). This finding suggests that human spatial navigation strategies are quite flexible, exploit multisensory information, and depend on the particular type of response that is required at the given moment. In contrast to previous desktop experiments asking for a simple homing response that demonstrated a preference for distinct reference frames ***(Gramann et al., 2005**,**2010;Lin et al., 2015)*** the current task showed that the majority of participants preferred an allocentric reference frame. Having to constantly update one’s own position as well as the position of other entities in space (landmark, home) likely fosters the use of an allocentric reference frame.

Moreover, the behavioral results in the landmark pointing task (figure 2A) might follow the leaky integrator model ***(Lappe et al., 2007;Harris and Wolbers, 2012)***. This model assumes that the encoded orientation, as the variable state, is incremented with movement by multiplying the actual orientation gain factor. This process tends to decay by an orientation-dependent leaky factor. In our trials, there were no visible landmarks within the navigation task. Therefore, participants could not use external landmarks to anchor their cognitive map. Instead, they had to derive their orientation with each turn based on idiothetic information only. Thus, errors accumulated at each turning point (figure 2A) and increased somewhat proportionally to the number of turns in the scenario. In other words, the errors in the landmark pointing task (at R1, R2a, and R5a, figure 1) are correlated to the number of turning points (figure 2A).

Using a spatial filter approach and subsequent clustering of independent components, we demonstrate the RSC to reflect specific aspects of the navigation task. EEG contains subcortical activity and allows to localize deeper brain structures ***(Seeber et al., 2019)***. Previous desktop studies have already revealed navigation-related activity in or near the RSC ***(Gramann et al., 2010;Chiu et al., 2012;Lin et al., 2015)***. However, even though theta oscillation is an important mechanism for computing head orientation and providing a grid cell network ***(Brandon et al., 2011;Koenig et al., 2011;Winter et al., 2015)***, it has not been reported in human brain imaging studies using stationary setups. Notably, we found a strong theta synchronization in the RSC during periods of heading changes, which indicates that physical rotations induce RSC activity (figure 3). This has not been reported in previous stationary studies. This theta oscillation was stably observed with each turn by navigators along the path. Through MoBI, our participants were able to make use of naturally occurring spatial information, such as motor efference and cues from the visual, vestibular, proprioception, and kinesthetic systems. Therefore, participants could extract their head direction from HD cells activity ***(Shine et al., 2016;Chen et al., 2018)***, which is often eliminated in stationary setups. In addition, there is evidence that the firing rate of HD cells decreases with restraints during navigation. Compared with those in active locomotion, loosely restrained rats in passive movement showed a nearly 24%reduction in the peak firing rates of HD cells ***(Zugaro et al., 2001;Bassett et al., 2005)***, while tightly restrained rats ***(Taube, 1995;Knierim et al., 1995)*** showed near or complete suppression. The suppression of the HD cell firing rate is due to disruption of the vestibular system, which is the essential signal for estimating head direction ***(Cullen and Taube, 2017;Stackman et al., 2003;Muir et al., 2009)***. In previous human spatial navigation studies, the population was limited to stationary participants. Thus, the vestibular information for HD signals may have been eliminated ***(Gramann, 2013)***. In this study, the participants received vestibular information in addition to all other naturally occurring sensory modalities while turning and walking. Accordingly, sufficient multi-modal sensory information was available for them to compute their head directions. Therefore, theta oscillation occurred after each turning onset (figure 3), indicating heading computation activity in the RSC, providing evidence of heading computation in healthy participants in a physical spatial locomotion study replicating similar theta activity in participants rotating on the spot ***(Gramann et al., 2018)***. Furthermore, we replicated RSC alpha suppression, which has been previously observed in spatial learning for maintaining orientation in both passive ***(Gramann et al., 2010;Plank et al., 2010;Chiu et al., 2012;Lin et al., 2015)*** and active navigation tasks ***(Ehinger et al., 2014)***. This alpha desynchronization might indicate ongoing spatial transformations from egocentric to allocentric coordinates and vice versa ***(Gramann et al., 2010)***. Although ***Kim and Maguire (2019)*** demonstrated that RSC activity is correlated with behavioral performance in three-dimensional space, how the use of distinct reference frames during navigation impacts RSC activity was still unclear. Remarkably, in this study, we found that RSC activity systematically covaried with behavioral responses, and that this correlation depended on the reference frame used. The use of an allocentric strategy revealed a positive correlation of individual performance and alpha power, while an egocentric strategy was associated with a negative correlation (figure 4). This general pattern was confirmed using a single-trial analysis approach that identified the reference frame underlying the single-trial pointing response of participants irrespective of their general reference frame proclivity. The results clearly indicate that only the use of an allocentric response and not the use of an egocentric response was associated with desynchronization in alpha band activity.

Overall, we exposed some of the behaviors and neural dynamics associated with physical navigation. We found that theta oscillations originating from the RSC are present during human navigation, more prominent while computing direction changes, and consistently synchronized irrespective of the length of the total path walked. Further, the results suggested that alpha desynchronization is essential for translating between spatial reference in active and passive navigation.As the first evidence of how our sense of direction works in healthy moving humans, these findings demonstrate that, for some, our brain dynamics are not the same when simulating navigating through space as opposed to actually moving through space.

## Acknowledgments

We thank H.T. Chen and A.K. Singh for their useful discussions, C.A. Tirado for help with the virtual reality programming, and all the participants for giving their time for this study. This work was supported in part by: the Australian Research Council (ARC) under discovery grant DP180100670 and DP180100656; the Australia Defence Innovation Hub under Contract No. P18-650825; and the US Office of Naval Research Global under Cooperative Agreement Number ONRG - NICOP - N62909-19-1-2058. We would also like to thank the NSW Defence Innovation Network and the NSW State Government of Australia for financial support for part of this research through grant DINPP2019 S1-03/09.

## Author contributions

Do, Gramann and Lin designed the experiment. Do acquired and analyzed the data together with Gramann. Do, Lin and Gramann wrote the manuscript. All authors discussed the results and contributed edits to the manuscript.

---

**Figure 3–Figure supplement 1.**
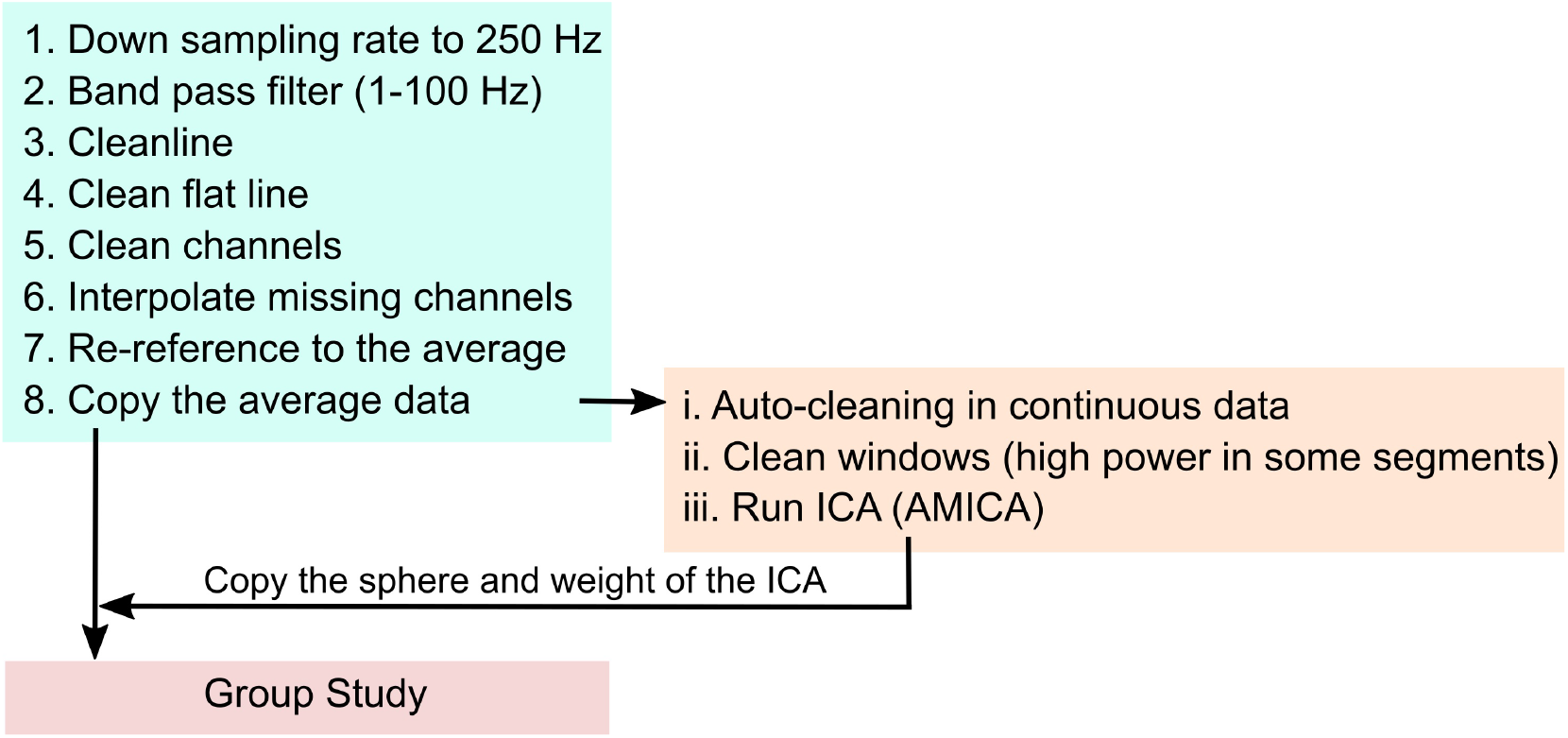
EEG pre-processing pipeline.

**Figure 3–Figure supplement 2.**
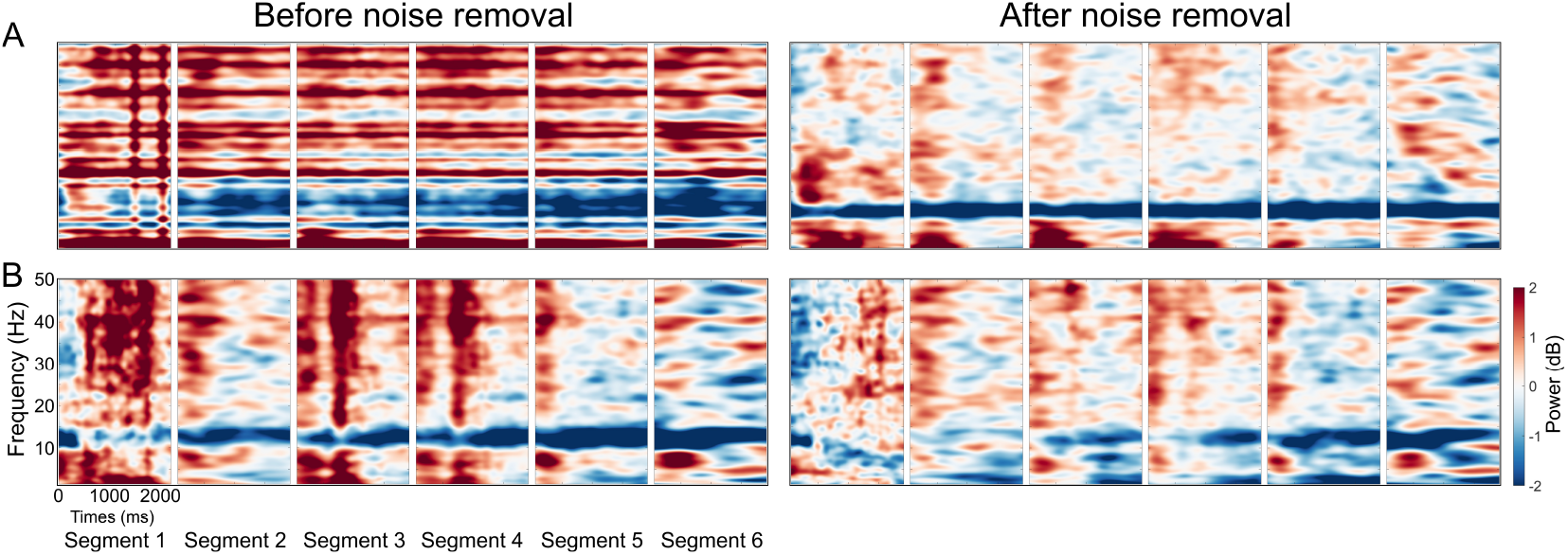
The RSP ERSP before and after noise removal at the participant level. The left figure shows the average ERSP for participants before noise removal. The right figure shows the average ERSP for participants after noise removal at: (**A**) one egocentric participant;and (**B**) one allocentric participant.

**Figure 3–Figure supplement 3.**
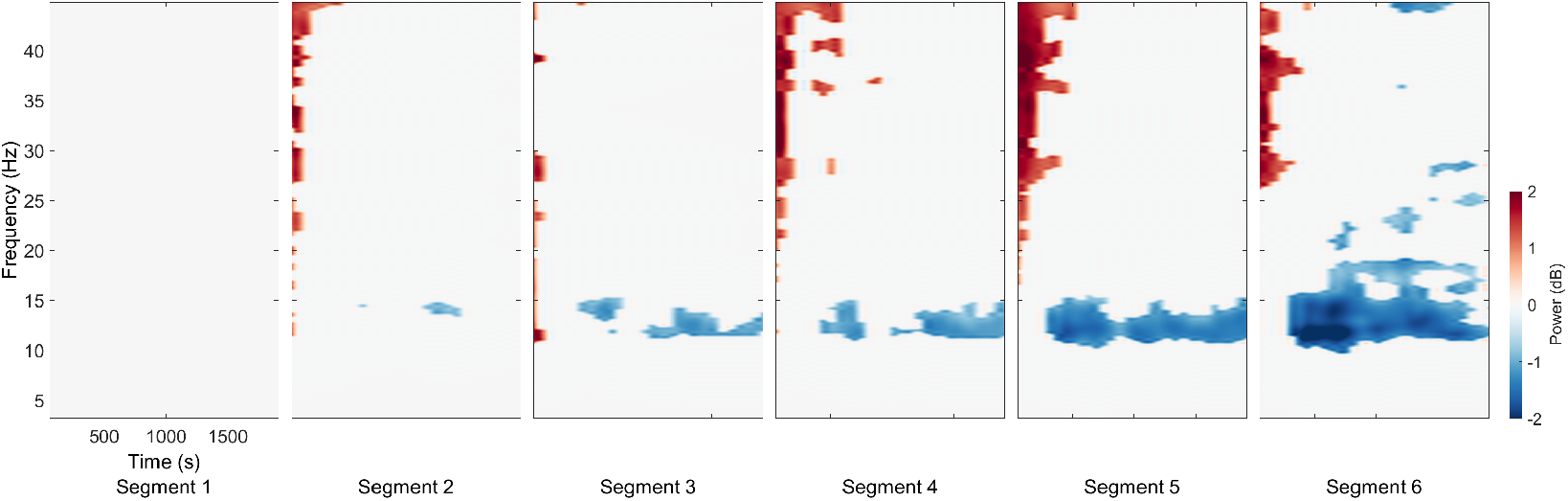
The permutation test (n=2000, FDR-correlated) of the RSC ERSP for 6 segments in comparison to segment 1.

**Figure 3–Figure supplement 4.**
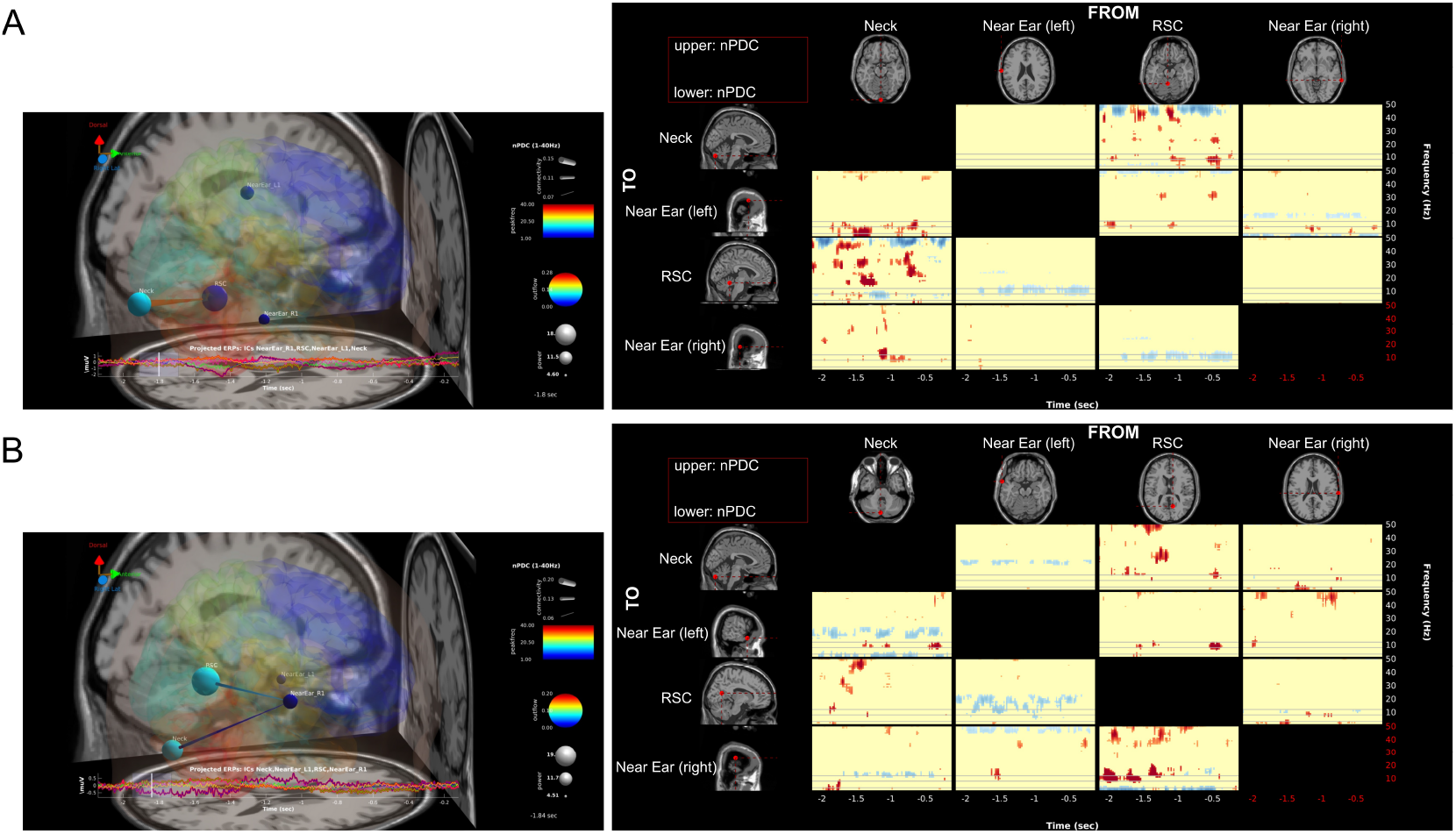
Effective connectivity. The estimated effective connectivity of the four clusters (the RSC, neck, near the ear on the left side, and near the ear on the right side) in one participant in the (**A**) allocentric and (**B**) egocentric strategy groups. The results indicate that theta activity in the neck cluster has no effect on the RSC.

**Figure 3–Figure supplement 5.**
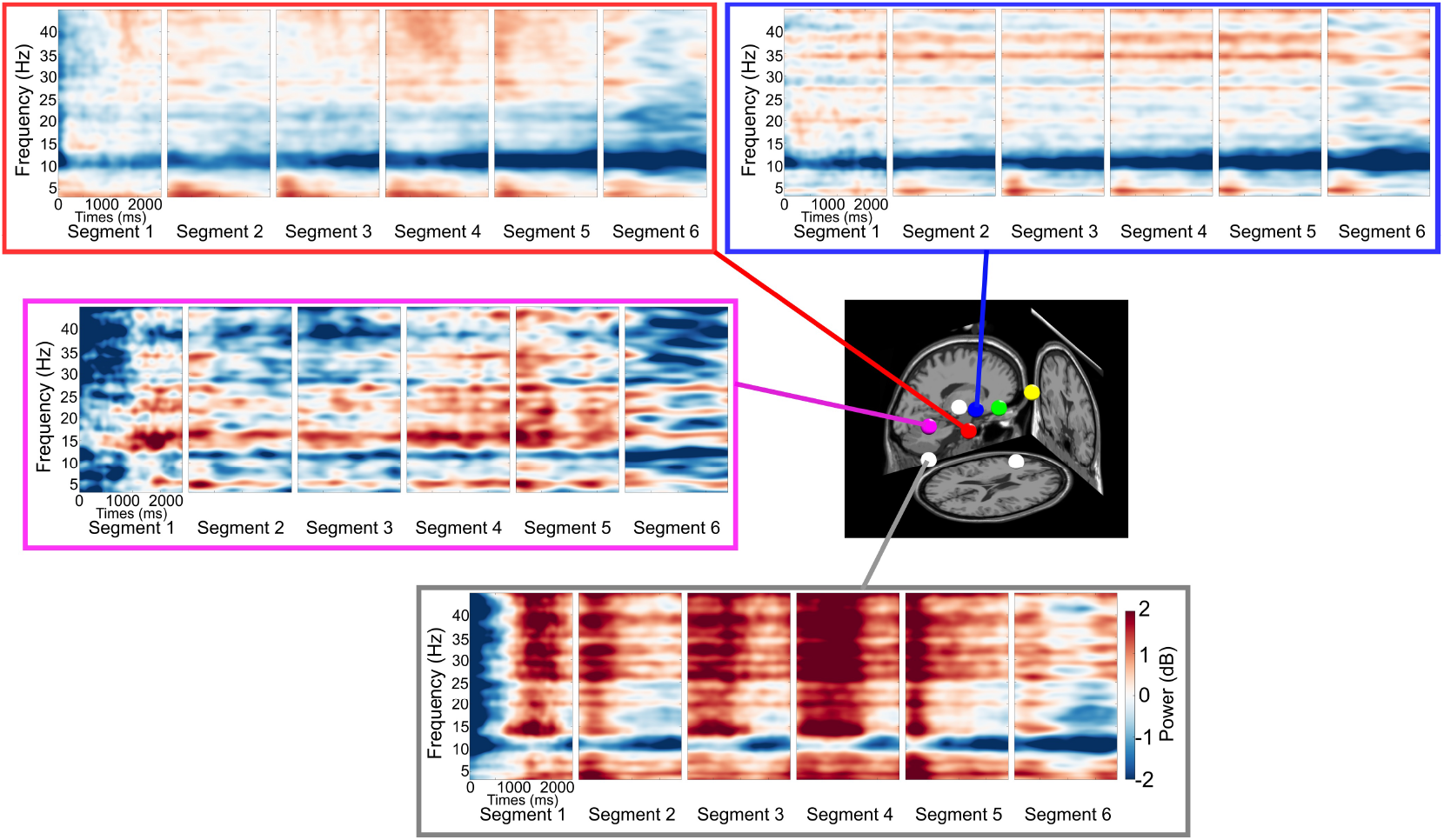
The ERSP in different brain regions in or near the RSC (red), parietal cortex (blue), occipital cortex (pink), and neck (white).

**Figure 4–Figure supplement 1.**
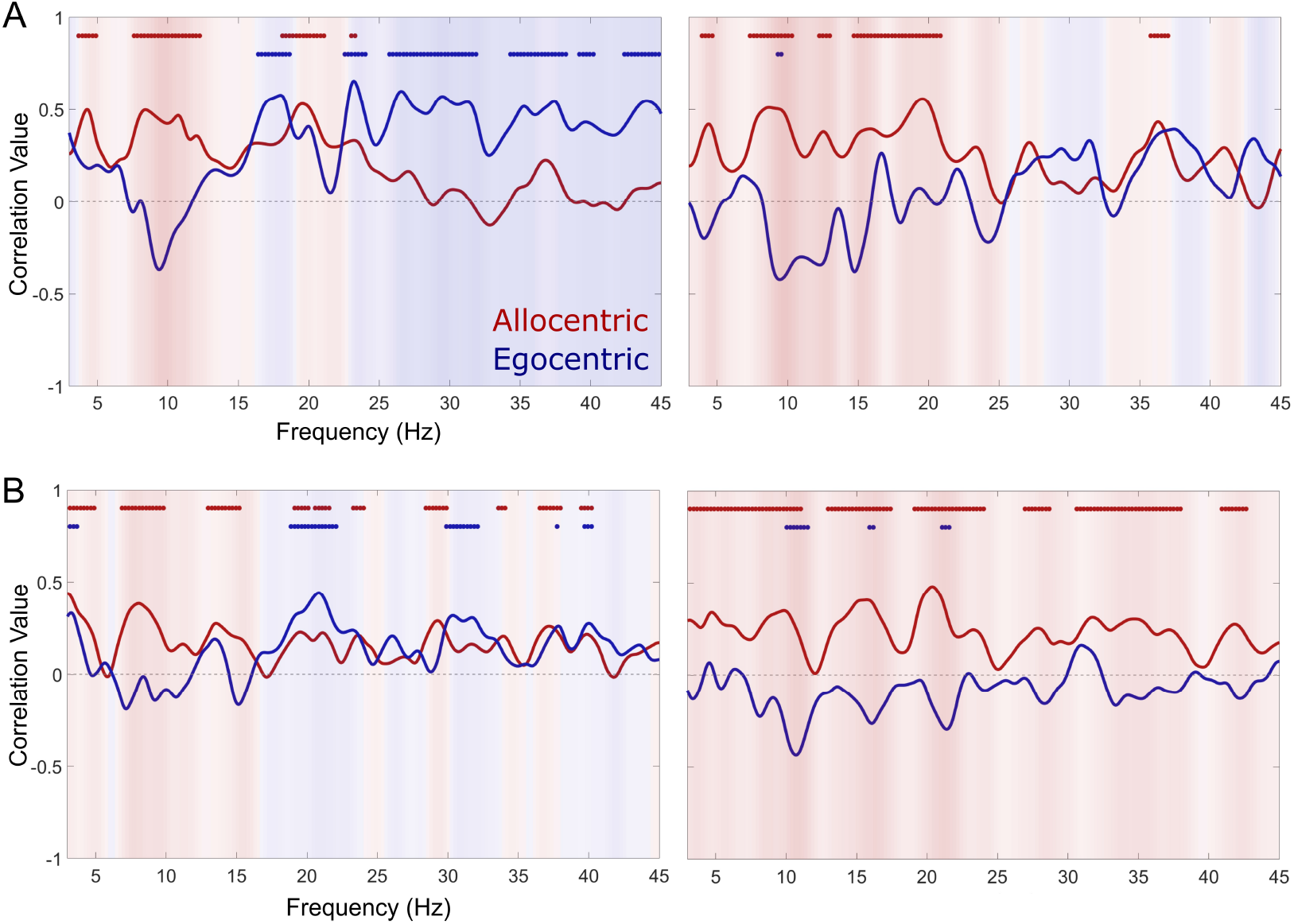
Correlations between pointing task errors and RSC power at: (**A**) the participant level;and (**B**) the trial level. These are correlation coefficients between performance and the RSC ERSP in the continuous frequency (3-45 Hz) in the first ten percent (left column) and middle ten percent (right column) of the segment length. The asterisk (*) indicates a significant difference (p<0.05) between the allocentric (red) and egocentric strategies (blue).

